# Cystine/glutamate antiporter system Xc^-^ deficiency impairs insulin secretion

**DOI:** 10.1101/2023.01.19.524735

**Authors:** Axel de Baat, Daniel T Meier, Leila Rachid, Adriano Fontana, Marianne Böni-Schnetzler, Marc Y Donath

**Author notes:** Corresponding author: Axel de Baat.

## Abstract

System Xc^-^, encoded by *Slc7a11*, is an antiporter that exports glutamate and imports cystine. Cystine is used for protein synthesis and incorporation in thiol peptides such as glutathione, which function as cofactors for reactive oxygen species scavenging enzymes. Glutamate export by astrocytes through system Xc^-^ has been implicated in excitotoxicity, a form of neurotoxicity that has been postulated to also occur in insulin-producing beta-cells in the pancreatic islets. This study describes the implications of *Slc7a11* deficiency on glucose metabolism in both constitutive and myeloid cells-specific knockout mice. Constitutive *Slc7a11* deficiency leads to drastically lowered glutathione levels in the pancreatic islets and immune cells in addition to diminished insulin secretion both *in vitro* and *in vivo*. Macrophage-specific deletion did not have a significant impact on metabolism or islet function. These findings suggest that system Xc^-^ is required for glutathione maintenance and insulin production in beta-cells, but is dispensable for islet macrophage function.

## Introduction

Insulin secretion of pancreatic beta-cells is coupled to the many by-products of beta-cell glucose catabolism and subsequent oxidative phosphorylation, ranging from ATP and glutamate(1) to reactive oxygen species (ROS)(2). This relatively unrestricted flux of glucose metabolism therefore makes beta-cells especially sensitive to redox stress induced by hyperglycemia(3)(4). One of the main drivers of Type 2 Diabetes-associated cellular dysfunction is excessive production and defective detoxification of ROS(5). Detoxification of ROS is performed, among others, by glutathione peroxidases that require the thiol-containing tripeptide glutathione as a co-factor. The limiting substrate for the synthesis of glutathione is cysteine, a semi-essential amino acid containing a thiol group that is useful for redox chemistry. While cysteine can be taken up by a range of amino acid transporters, its oxidized dimeric form cystine is mainly taken up through system Xc^-^ (sXc). SXc is a heteromeric amino acid transporter comprised of a heavy chain (CD98), that it shares with many other amino acid transporters, and a unique light chain encoded by *Slc7a11*. The transporter functions as an anti-porter exchanging cystine for glutamate in a 1:1 ratio in a sodium-independent fashion(6). *Slc7a11* is upregulated upon cystine starvation by the activating transcription factor 4 (Atf4), in a mechanism where *Atf4* is preferentially transcribed when regular transcription slows down(7). When ROS accumulate within the cell, cysteine residues on nuclear factor erythroid-2 related factor 2 (NRF2) are oxidized. NRF2 then dissociates with KEAP1 resulting in nuclear translocation, which activates a network of genes involved in redox maintenance, including *Slc7a11*(8). In hypoxic conditions and inflammation, the breakdown of hypoxia-inducible factor 1-alpha (HIF1-a) is interrupted, leading to nuclear translocation and induction of *Slc7a11* transcription(9). SXc inhibition with Erastin leads to a type iron-mediated cell death called ferroptosis, a process that has also been suggested to occur and impair islet viability during transplantations[10]. Additionally, sXc in macrophages is also involved in diabetic retinopathy and wound healing[11][12]. When cells are low on cysteine, glutathione-specific gamma-glutamylcyclotransferase 1 (encoded by *Chac1*) is upregulated. CHAC1 degrades glutathione, thereby releasing cysteine from the tripeptide. This allows the cell to prioritize protein synthesis over maintaining a large glutathione pool, suggesting that glutathione is not only used for redox maintenance and detoxification but also as a store of cysteine(12).

Obesity is associated with macrophage infiltration and activation in tissues(13), during which profound metabolic and transcriptomic changes within the cell drive an anabolic response(14). This response includes upregulation of *Slc7a11* to supply enough cysteine for the inflammatory response and to mitigate the associated increased ROS production. These macrophages then produce and secrete pro-inflammatory cytokines such as IL-1 beta in order to re-establish homeostasis(15). When homeostasis cannot be fully restored, low-grade inflammation emerges that is characteristic of chronic metabolic disease(16).

Glutamate export by myeloid cells through sXc has been implicated as an important factor in excitotoxicity, a neuronal cell-death that is characterized by excessive glutamate-induced depolarization(17) (18). Beta-cells and neurons share developmental transcription factors, evolutionary origins and the utilization of membrane depolarization to release secretory granules(19). It has previously been described that beta-cells are sensitive to glutamate-induced cytotoxicity(9)(20). Given the high sensitivity of beta-cells to redox stress and the possible occurrence of excitotoxicity, we hypothesized that myeloid sXc activity plays a critical role in this process. To test this hypothesis, we used constitutive and conditional genetic models of *Slc7a11* deficiency and studied consequences on metabolism and macrophage physiology *in vivo* and *in vitro*.

## Materials and methods

### Mouse Models

All animal experiments were conducted according to the Swiss Veterinary Law and Institutional Guidelines and were approved by the Swiss Authorities. High fat diet was lard-based and obtained from Ssniff (Soest, Germany).

Constitutive Slc7a11 knockout mice were first generated by Sato et al.(21). The transgene was propagated by crossing heterozygous parents in order to ensure littermate representation in both experimental groups. C57Bl/6N Slc7a11<tm1a(EUCOMM)Wtsi/IcsOrl> were acquired from INFRAFRONTIER GmbH and imported to our facility by in vitro fertilization of acquired sperm. To generate the tm1b allele, mice were first crossed with B6-Tg(ACTFLPe)9205Dym/NPg to induce flipase-mediated recombination of FRT-flanked resistance cassette. To generate a conditional myeloid cells-specific Slc7a11 knockout, mice were crossed to B6N.129P2-Lyz2<tm1(cre)Ifo>, acquired from Jackson and backcrossed to C57Bl/6N background. The resulting mouse line did not exhibit any observable developmental defects. Knockdown efficiency was assessed by quantitative PCR.

### Endotoxemia

To induce endotoxemia, mice were injected with 2mg/kg body weight phenol-extracted LPS (Escherichia coli O26:B6, Sigma-Aldrich) dissolved in PBS. Mice were monitored regularly during the 2-8 hours before sacrifice or metabolic experiments.

### Intraperitoneal Glucose and Insulin tolerance tests

For the intraperitoneal glucose tolerance test, mice were fasted for 6 hours (from 8AM until 2PM) and then injected with 2g/kg body weight glucose with a 40% glucose solution. Blood glucose was monitored at the 0, 15, 30, 60, 90 and 120 minute timepoint post injection using a Freestyle Lite glucometer form Abbott. Additionally, at the 0, 15 and 30 minute timepoint, 25 uL of blood was collected in tubes containing 2.5 uL of 50mM EDTA solution to obtain plasma for insulin measurements.

Insulin resistance was assessed by insulin tolerance tests where mice were injected with 1 (1.5 for high-fat diet-fed mice) U/kg body weight Actrapid. Glucose was measured at the 0, 15, 30, 60, 90 and 120 minute timepoint post injection using a Freestyle Lite glucometer from Abbott.

Glycemic index was calculated as: (Insulin at t=15 - Insulin at t=0)/(Glycemia at t=15 - Glycemia at t=0)

### (3H)2-deoxyglucose uptake and detection

To assess tissue glucose uptake, mice were fasted for 6 hours and injected with 2g/kg bodyweight 40% glucose solution spiked with 350 uCi/kgbw (3H)2-deoxyglucose acquired from Perkin Elmers. Tissues were harvested and 200 mg of the respective tissue was taken and homogenized in 500 uL ultra-pure water with a stainless-steel bead in a TissueLyser II. The homogenates were then heated in a heating block at 95 °C for 5 minutes and subsequently pelleted. Supernatants were collected and phosphorylated 2-(3H) deoxyglucose was separated by Poly-Prep columns (BioRad). The final eluates were mixed with Ultima Gold Scintillation fluid at a 1:7 Ratio in scintillation tubes and analysed with a beta counter.

### Adipose tissue isolation

Epididymal white adipose tissue was isolated and minced with scissors. The adipose tissue was then incubated with collagenase for 45 minutes at 37 °C in a shaker. Stromal vascular fraction was obtained through centrifugation.

To isolate adipocytes, the digested adipose tissue was spun at 30 x g, the infranatant was removed, the floating fraction was washed and filtered through a 200 um pore size mesh. Adipocytes were then distributed for experiments based on volume.

### Ex-vivo adipocyte 2-DG uptake

To assess glucose uptake in isolated adipocytes, Promega glucose-uptake glo assay based on 2-Deoxy Glucose was used. In short, isolated adipocytes were distributed, normalized by volume, in glass tubes in a 37 °C shaker in modified Krebs-Ringer bicarbonate buffer (KRB; 115 mM NaCl, 4.7 mM KCl, 2.6 mM CaCl_2_ 2H_2_O, 1.2 mM KH_2_PO_4_, 1.2 mM MgSO_4_ 7H_2_O, 10 mM HEPES, 0.5% bovine serum albumin, pH 7.4) containing 11mM Glucose and 1mM 2-DG +/-100nM insulin. After shaking incubation for 1 hour, supernatant was removed and cells were washed twice with KRB containing 11mM Glucose. Cells were subsequently processed according to manufacturer’s protocol.

### Islet isolation

Islets were isolated by collagenase digestion through infusion of collagenase mix through biliopancreatic duct. Islets were harvested by handpicking under a microscope and cultured in a cell culture incubator (37 °C and 5% C02) in DMEM containing 11.1 mM glucose, 1mM Pyruvate, Non-Essential Amino Acids, 100 units/ml penicillin, 100 μg/ml streptomycin, 2 mM Glutamine, 50 μg/ml gentamycin, 10 μg/ml Fungison and 10% FCS.

### Glucose-stimulated insulin secretion

GSIS protocol in 96 well plates was adapted from an earlier publication(22). In short, islets were incubated in 96 well plates with 1 islet per well for 2-3 days in a cell culture incubator in DMEM. Islets were washed and then pre-incubated for 1 hour in preequilibrated modified Krebs-Ringer bicarbonate buffer containing 5.5 mM glucose. Followed by 1 hour incubation to obtain basal secretion, subsequently the medium is exchanged for 16.7mM glucose KRB, yielding the stimulated sample. After the experiment, wells were checked by eye and wells not containing an islet were excluded. Supernatants from five wells were pooled per sample and 3-4 samples were taken per mouse. Insulin was quantified with the mesoscale mouse/rat insulin assay. Stimulation index was calculated as the ratio of stimulated to basal secretion. γ-Glutamyl-cysteine was kindly provided by Martin Zarka from Biospecialities.

### RNA Isolation and quantitative real-time PCR

RNA was isolated by the Nucleo Spin RNA II Kit. cDNA was prepared with the GoScript Reverse Transcription Mix containing random primers according to the manufacturer’s instructions. For quantitative qPCR of *Chac1, Il1b* and *Slc7a11* SYBRgreen-based chemistry with GoTaq Polymerase and the ABI 7500 fast system was employed. Hprt was used as a housekeeping gene and expression levels were calculated as –(gene of interest – housekeeping gene) as to not assume an amplification efficiency of 2 and to display the data in the most unadulterated manner. Primer sequences were all derived from the first ranked pairs in the Harvard Primer Bank https://pga.mgh.harvard.edu/primerbank/.

### Flow cytometry

Flow cytometry experiments were performed on a CytoFLEX V2-B4-R2 Flow Cytometer (8 Detectors, 3 Lasers).

For measuring intracellular thiol levels, Thioltracker dye was used according to manufacturer’s directions. Cells were washed and subsequently analysed.

### FACS sorting of macrophages and dispersed islets fractions

After isolation, islets were dispersed and beta-cells were sorted based on size and granularity as described before(23) and islet immune cells were sorted as the CD45+ fraction.

Macrophages from different tissues were identified as F4/80 PE-Cy7 and CD11b PerCP-Cy7 positive cells after being gated for singlet, DAPI-negative, lineage negative (CD19, Ly-6G, CD3, NK1.1, BV510). The cells were sorted on the FACSMelody.

### ATP, Glutathione and redox potential measurements

ATP measurements were performed using CelltiterGlo3D. For glutathione measurements in islets, an adapted version of the glutathione-glo assay was performed where the standard lysis buffer was replaced with passive lysis buffer in order to prevent inadequate permeabilization. One sample represents 10 islets.

### Microscopy and beta-cell mass quantification

The weight of the pancreas was determined at sacrifice. Staining for insulin and CD45 were performed as previously described[25]. Pancreas and beta-cell area were determined using a convolutional neural network approach using the U-net architecture from https://github.com/jvanvugt/pytorch-unet. The outputs were hand curated and analysed using ImageJ software. Normalized beta-cell weight was calculated as ratio (beta-cell area/pancreas area)*(pancreas weight/bodyweight).

### Data analysis

Results were analyzed with Prism 9.1.0 (GraphPad, USA) or Python and Jupyter notebook (Jupyter Labs), p < 0.05 was considered to be significant. Results were expressed as mean ± SD. Data were analyzed with unpaired two-sided Mann–Whitney U test or 2-way ANOVA. Statistical details of experiments are described in the figure legends and in the figures.

## Results

### Slc7a11 deficient-mice have increased insulin secretion and glucose disposal that is abrogated upon aging and high-fat feeding

To understand the role of sXc in glucose metabolism, we obtained constitutive *Slc7a11* deficient mice, generated by Sato et al. through insertion of a selection cassette in exon 1(21). As previously described, no developmental alterations were observed in these mice. At 12 weeks of age, glucose disposal during an intra-peritoneal glucose tolerance test (ipGTT) was increased in knockout (KO) animals (fig 1a, b) along with increased insulin secretion (P=0.087 for insulinogenic index, fig 1c-e). Body weight was comparable between the groups (fig 1f).

**Figure 1.**
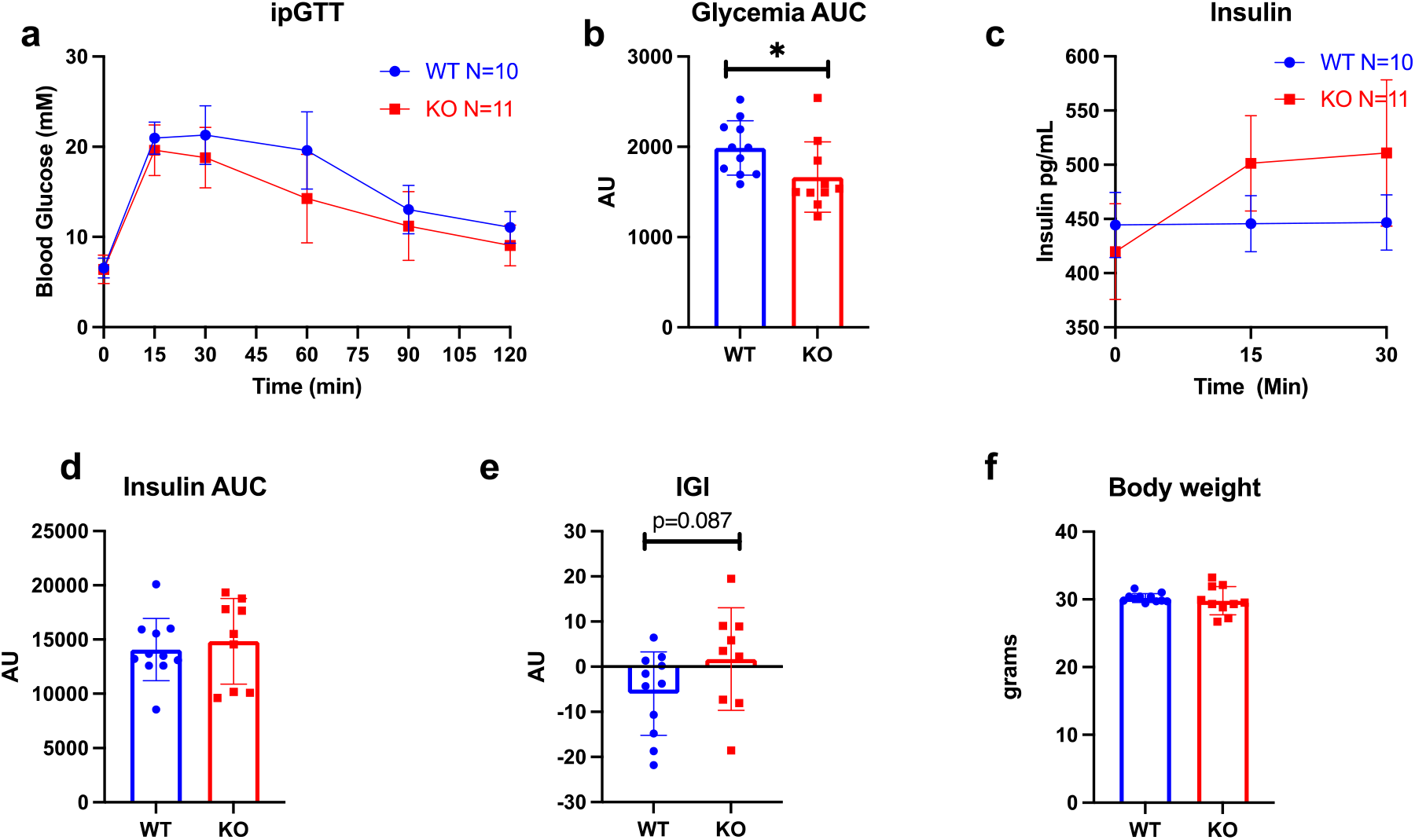
sXc Whole Body KO and Wild Type (WT) mice at 12 weeks of age (a) ipGTT glycemia, (b) corresponding AUC, (c) plasma insulin and (d) corresponding AUC after intraperitoneal injection of 2g/kg glucose following a 6h fast, (e) Insulinogenic Index, (f) body weight. Statistics: two-sided Mann–Whitney U test; error bars represent SD; ★p < 0.05.

At 26 weeks of age, increased glucose disposal was still apparent (fig 2a, b), although insulin secretion started to deteriorate compared to controls (fig 2c, d, e) without changes in body weight (fig 2f). Diminished insulin secretion was not driven by reduced beta-cell mass (fig 2g). Glucose uptake after a glucose bolus was increased in the epidydimal adipose tissue and muscle (fig 2h, i). In adipocytes isolated from sXc knockout mice, basal glucose uptake was increased but failed to be further enhanced in response to insulin (fig 2j). These data suggest that glucose metabolism is improved in sXc knockout mice due to increased insulin secretion at younger age which transitions to lower insulin secretion in later life that is compensated by increased glucose disposal.

**Figure 2.**
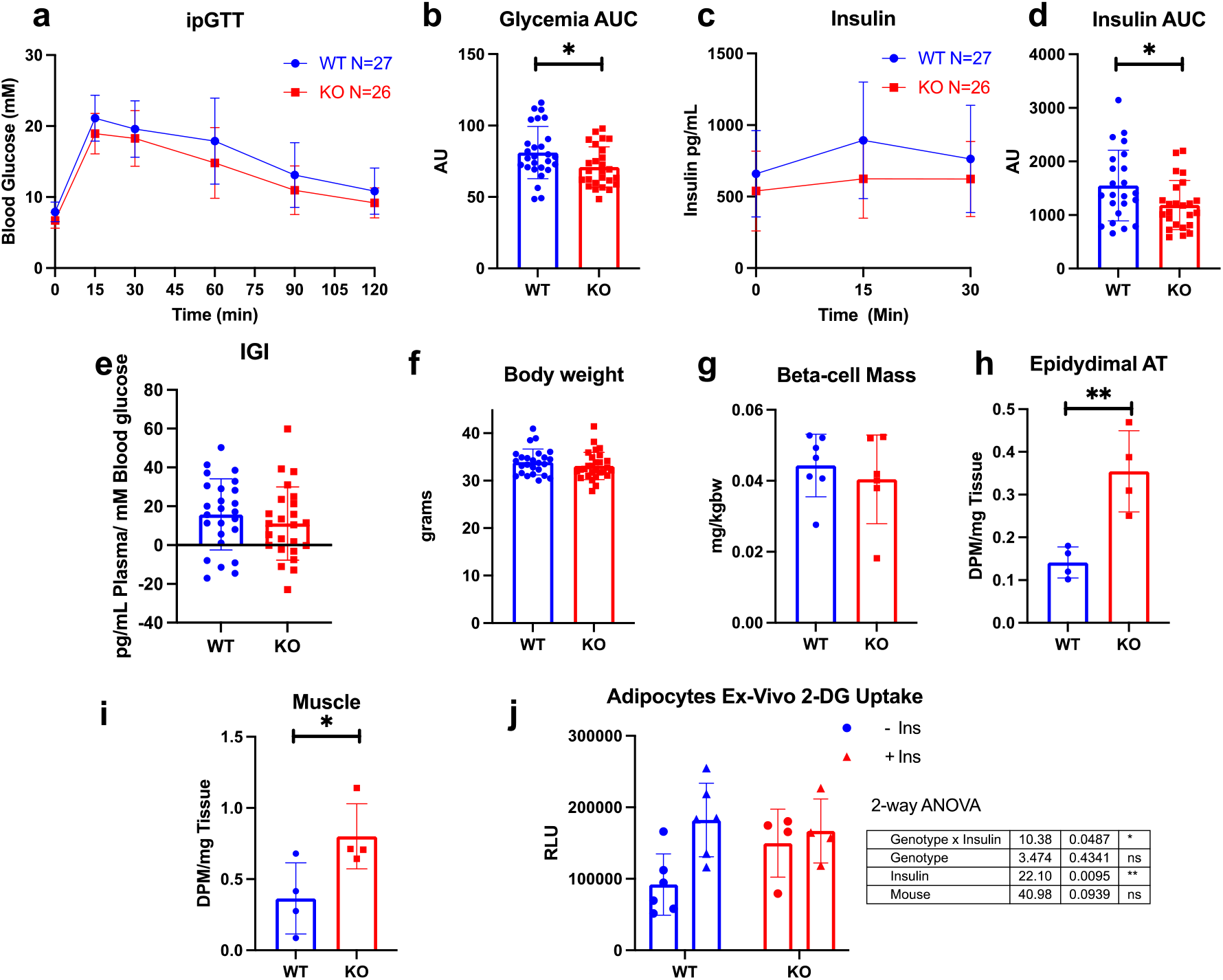
Slc7a11 deficient mice at 26 weeks of age fed normal chow have increased glucose disposal and lower insulin secretion (a) ipGTT glycemia, (b) corresponding AUC, (c) plasma insulin and (d) corresponding AUC after intraperitoneal injection of 2g/kg glucose following a 6h fast. (e) Insulinogenic index (f) Body weight, (g) beta-cell mass estimated by histology histology and normalized to pancreas and body weight, [3H]2-deoxyglucose uptake in (h) Epididymal Adipose Tissue and (i) Muscle, (j) glucose uptake in isolated adipocytes in response to 100nM insulin, each datapoint represents 1 animal. Analysed with 2 way ANOVA, Statistics: two-sided Mann–Whitney U test, 2-way ANOVA; error bars represent SEM; ★p < 0.05, ★★p < 0.01.

Next, we investigated the implications of sXc-deficiency upon a high calorie diet. 8-week-old mice were fed a high-fat diet (HFD) for 4 weeks. In contrast to mice on a chow diet, glucose disposal was not significantly different between genotypes (fig 3a, b). Similar to aged mice, insulin secretion was reduced in KO animals (fig 3c, d), while insulinogenic index was unchanged (fig 3e). There was no difference in body weight (fig 3f), beta-cell mass (fig 3g) or glucose uptake in epidydimal adipose tissue (fig 3h). Finally, we analysed the expression of *Chac1* in islets as an indication of cysteine shortage. Indeed, *Chac1* expression levels in FACS-sorted beta-cells and islet immune cells were increased (fig 3i). These experiments suggest that sXc knockout modestly increases insulin secretion and enhances glucose disposal in mice on a chow diet initially, while these effects disappear with age or during obesity.

**Figure 3.**
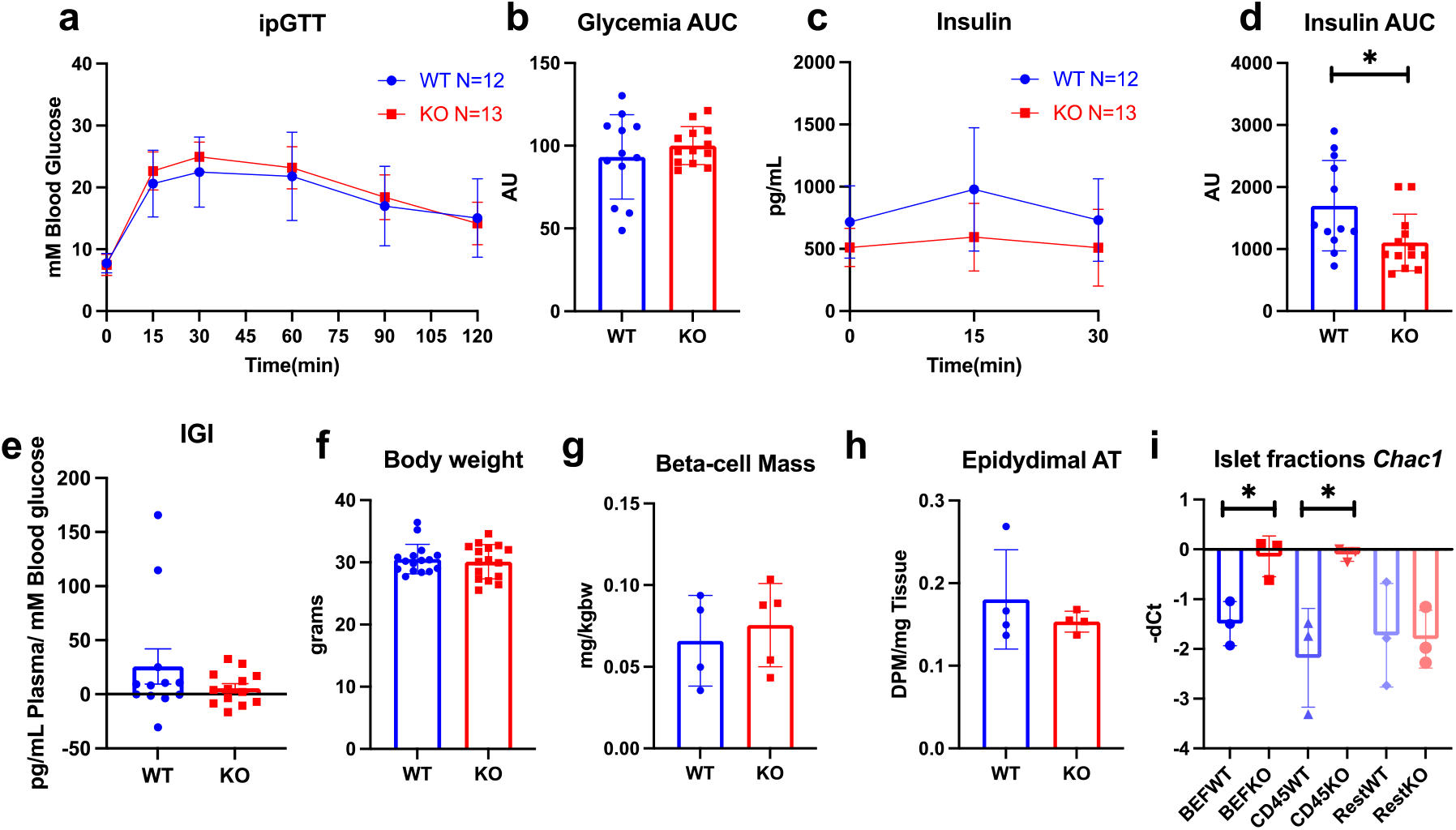
12-week-old whole-body male Slc7a11 deficient mice after 4 week HFD (a) ipGTT glycemia, (b) corresponding AUC, (c) plasma insulin and (d) corresponding AUC after intraperitoneal injection of 2g/kg glucose following a 6h fast, (e) Insulinogenic index(f) body weight, (g) Beta-cell mass determined by histology, (h) 3(H)2-deoxyglucose uptake in Epididymal Adipose Tissue (i) Chac1 expression in FACS isolated islet fractions, BEF= Beta-cell enriched, CD45 = CD45 positive fraction, Rest= Low sidescatter single cells. Statistics: two-sided Mann–Whitney U test; (a,b) error bars represent SD; ★p < 0.05, ★★p < 0.01.

### SXc is required to maintain islet glutathione levels and glucose-stimulated insulin secretion

To support the role of sXc in insulin secretion, we used the sXc inhibitor Erastin. Exposure of islets isolated from wild-type mice to Erastin for 24h lowered glucose-stimulated insulin release (fig 4a, b). When Erastin was added only to the glucose-stimulation medium, insulin secretion was almost completely abolished (fig 4a, b). Further, islets isolated from xCT knockout mice also showed diminished glucose-stimulated insulin secretion at both basal and glucose-stimulated conditions (fig 4c-d). We then wondered if supplying glutathione would rescue the impaired secretory phenotype. Pre-treatment of islets isolated from Slc7a11 knockout mice and controls with the glutathione precursor γ-glutamyl-cysteine removed the previously observed difference between the groups. The treatment also generally reduced the amount of insulin secreted in islets from both genotypes however (fig 4c,d).

**Figure 4.**
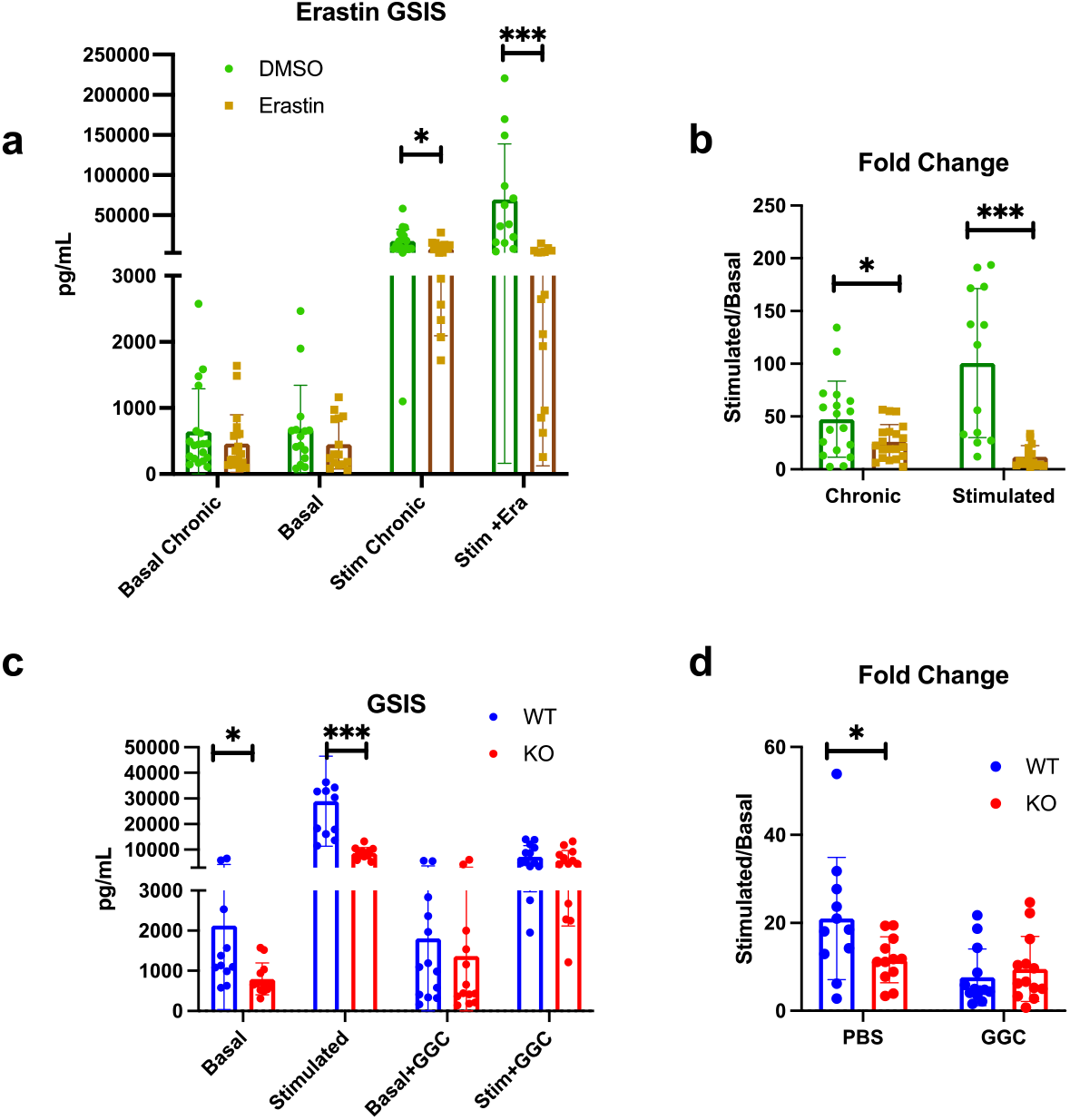
Effect of the sXc inhibitor Erastin (5 uM) on insulin secretion in Basal= 5.5mM and Stim 16.7 mM glucose Krebs ringer buffer (a-b) GSIS of islets treated with 5uM Erastin for 24 hours, GSIS of islets exposed to 5uM Erastin in the high glucose (stimulated) medium. (c,d) Glucose stimulated insulin secretion islets WT and KO islets including treatment with 1mM γ-glutamyl-cysteine for 24 hours, each dot represents a pooled sample of 5 islets, 3 mice were used per group Statistics: two-sided Mann–Whitney U test; error bars represent SD; ★p < 0.05, ★★★p < 0.001. Each dot represents a pooled sample of 5 islets, islets from 3 different mice were used per experiment, 3 experiments in total.

Next, we analysed ATP levels in isolated islets and found a reduction in ATP content of islets from sXc KO mice compared to islets from WT littermate controls (fig 5a). We then determined glutathione levels in whole islets and found it to be significantly lower in islets from KO mice (fig 5b). γ-Glutamyl-cysteine treatment partially prevented this reduction of glutathione in KO islets but also reduced glutathione levels in WT islets to a level comparable to KO islets given the same treatment. An additional group of islets was treated with Erastin, which also lowered the glutathione levels in KO islets treated with γ-Glutamyl-cysteine. This was surprising since the islets lack sXc, and suggests non-sXc-mediated effects on glutathione levels by Erastin. Consistent with lower glutathione levels, expression of *Chac1* was upregulated in FACS-sorted beta-cells isolated from 4 week HFD-fed animals. *Atf4*, a gene encoding a transcription factor that mediates a stress response upon halting of eukaryotic Initiation Factor mediated transcription by for example cysteine shortage, did not differ between genotypes (fig 5c). Given the lower concentrations of the redox enzyme co-factor glutathione, we next measured ROS and lipid peroxidation. Surprisingly, there was no difference between WT and KO islet fractions, suggesting that the lower glutathione levels are still adequate for maintaining redox homeostasis (fig 5d). These data support the hypothesis that ablation of sXc lowers insulin secretion which is likely not driven by excessive redox stress but could be the result of a mild cysteine shortage that further diminishes glutathione levels through *Chac1*. Finally, we evaluated whether the lower glutathione levels result in lower thiol levels and found thiols in beta-cells to be similar between wild-type, sXc deficient or wild-type beta-cells treated with Erastin (Fig 5e). However, SXc inhibition reduced thiol levels in the islet immune cell fraction (fig 5e).

**Figure 5.**
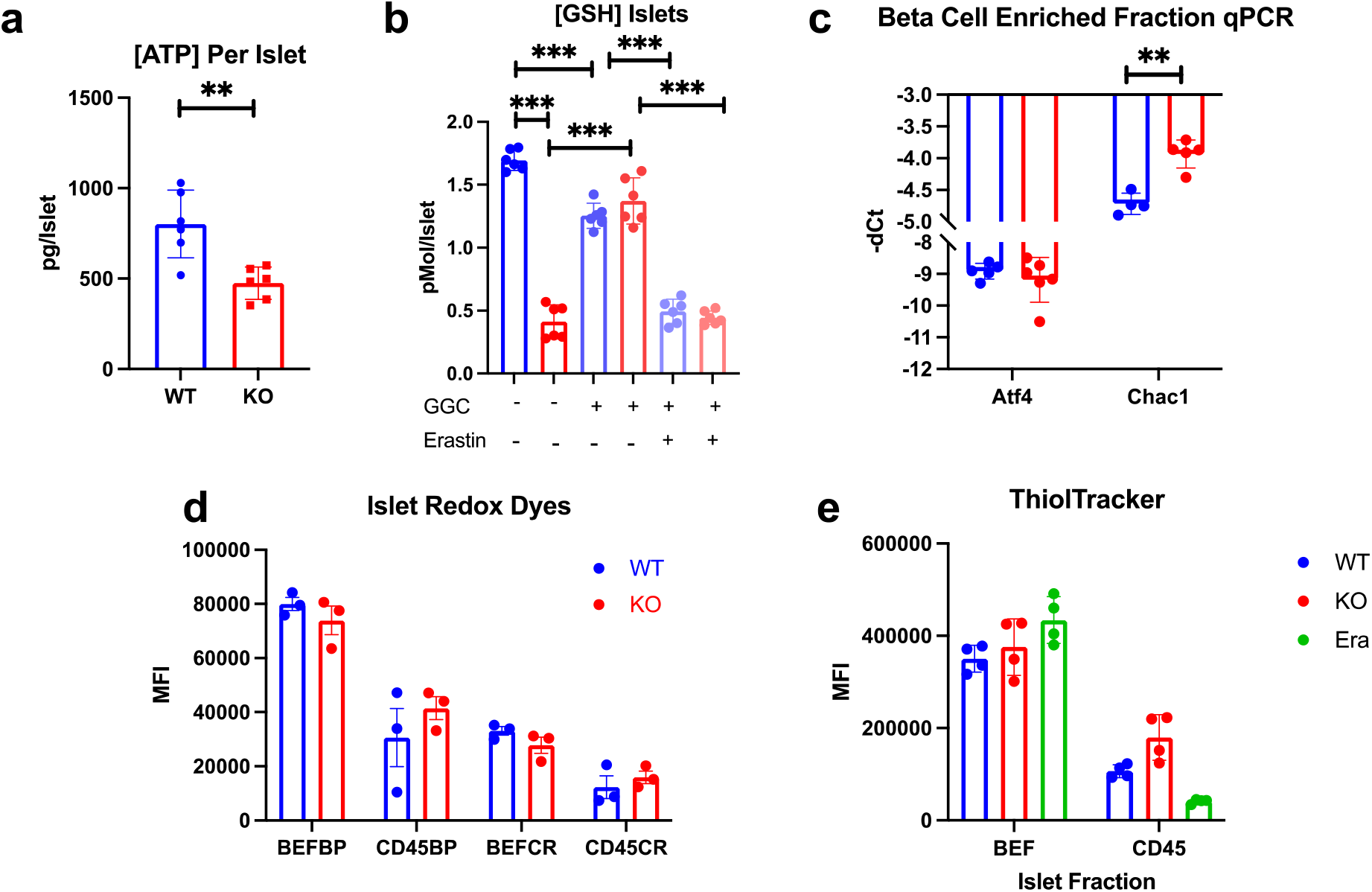
Functional aspects of isolated islets from 16-20 week-old male xCT knockout mice. (a) ATP levels per islet, (b) Glutathione concentration in whole islets, (c) qPCR of FACS sorted beta-cell enriched fraction, (d) ROS and Lipid peroxidation measured by flow cytometry with Cellrox green (CR) and Bodipy C11 (BP) MFI = mean fluorescence intensity, (e) Thioltracker thiol staining in dispersed islet fractions, BEF = Beta-cell enriched fraction, CD45 = CD45 positive fraction. (a,b) Each dot represents a 10 islet sample from 1 mouse, (c,d,e) each dot represents one mouse. Statistics: two-sided Mann–Whitney U test; error bars represent SD; ★p < 0.05, ★★★p < 0.001.

### *In vivo* LPS treatment does not lead to a differential response when comparing sXc deficient and WT animals

Because lipopolysaccharide (LPS) exposure leads to rapid upregulation of *Slc7a11* [19], we hypothesized that LPS pre-treatment might emphasize sXc’s role in metabolism. We therefore injected whole-body sXc knockout mice with 2mg/kgbw LPS 6 hours before an ipGTT. We confirmed the previously reported increase in insulin secretion and glucose disposal(24) but failed to detect a differential response between WT and KO animals (fig S1a-c). Additionally, plasma IL-1beta levels were similar between genotypes (fig S1d). Whole blood glutathione levels trended towards a decrease in KO animals, but showed no differential response to LPS administration (fig S1e). These data show that sXc-deficiency does not have overt implications *in vivo* for cytokine production, mitochondrial membrane potential, and glucose metabolism.

### Myeloid-specific sXc deletion does not phenocopy whole-body KO mice

Macrophages within islets have previously been shown to be involved in the regulation of insulin secretion [27]. Furthermore, *Slc7a11* expression levels in the immune cell fraction was 225-fold higher compared to other islet cell fractions (fig 6a). We therefore crossed mice carrying the *Slc7a11* gene with exon 3 flanked by loxP sites to *Lyz2*-Cre mice to obtain myeloid cell-specific *Slc7a11* knockout mice (fig S2). Slc7a11 expression was reduced by 83 +/-9 % (SD) in sorted islet immune cells, showing efficient gene knockdown in this model (fig 6a). The improved metabolic phenotype observed in whole-body xCT KO mice at 26 weeks of age was not present in mice with a myeloid-specific deletion (fig 6b,c). Similarly, *ex vivo* we did not find a significant difference in insulin secretion in these islets (fig 6d). In contrast to our findings in the whole-body KO, the myeloid cell-specific knockout did not display a decrease in islet-wide glutathione levels (fig 6e). We fed myeloid sXc deficient mice a HFD for 4 weeks. Glycemia following a glucose-bolus was unchanged between the groups but insulin secretion trended lower in Cre-carrying animals, although this did not reach statistical significance (fig 6f,g). Body weight, beta-cell mass and *Chac1* expression in the beta-cell-enriched fraction was similar between genotypes (fig 6h-j). In sorted adipose tissue and peritoneal macrophages from *Lyz2*-Cre *Slc7a11* KO mice there was a trend towards higher *Chac1* expression levels relative to controls (fig S3c). Further, female *Lyz2*-Cre *Slc7a11* KO mice fed a HFD for 4 or 12 weeks also did not show altered insulin sensitivity or glucose metabolism (fig S4). We additionally performed glucose tolerance tests with animals fed HFD for 12 and 44 weeks and did not find a difference (fig S5). These data suggest that despite a high expression level of sXc in islet macrophages relative to endocrine and other islet cells, constitutive myeloid cell-specific sXc deficiency does not affect islet function.

**Figure 6.**
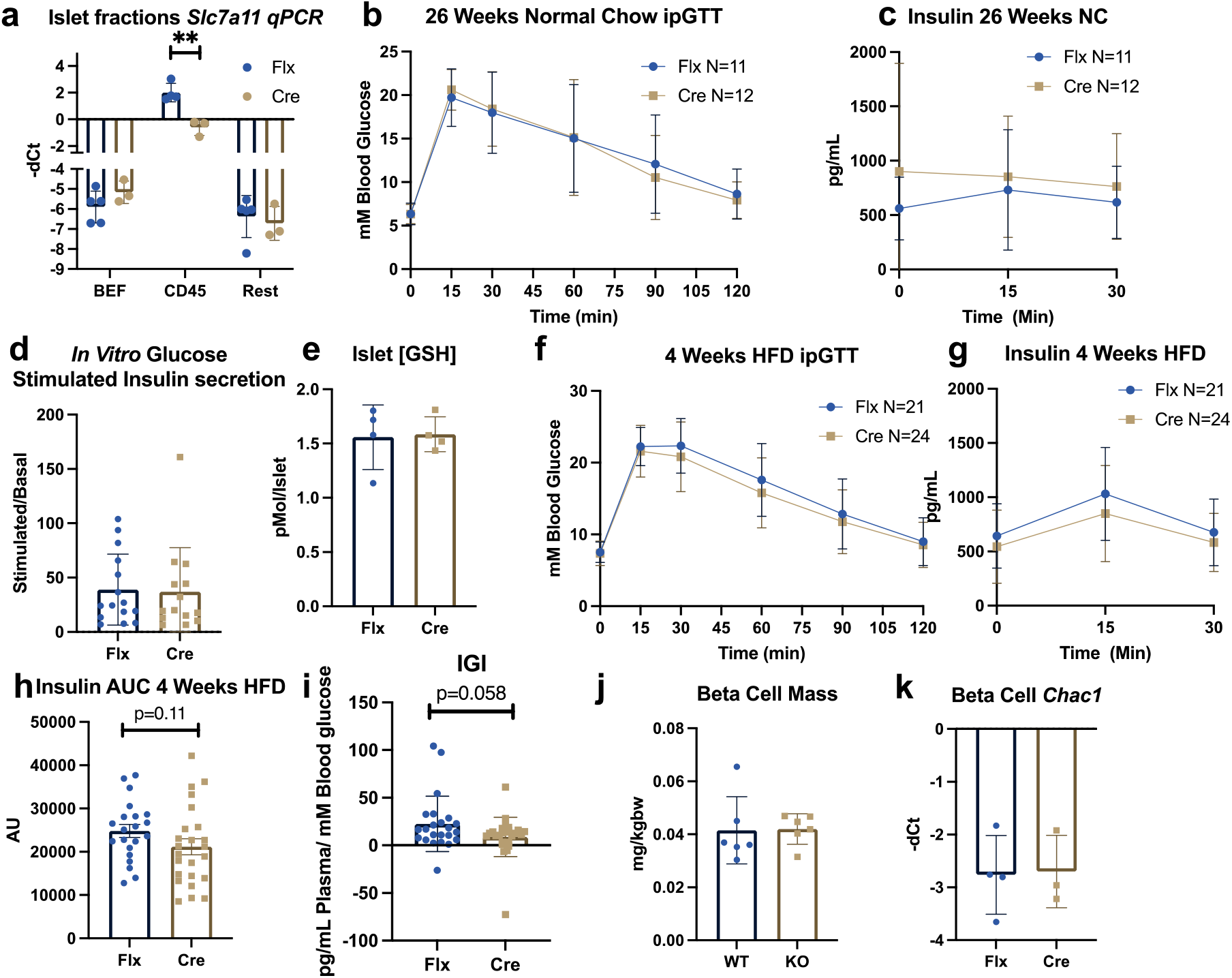
26-week-old LysM Cre Slc7a11 Flx (a) qPCR for expression of *Slc7a11* in sorted islet fractions, (b) ipGTT glycemia, (c) plasma Insulin after intraperitoneal injection of 2g/kg glucose following a 6h fast, (d) GSIS of isolated islets, (e) glutathione levels in whole islets normalized per islet. LysM Cre Slc7a11 Flx fed HFD for 4 weeks (f) ipGTT glycemia (g) plasma Insulin and (h) corresponding AUC after intraperitoneal injection of 2g/kg glucose following a 6h fast, (i) insulinogenic index, (j) beta-cell mass, (k) qPCR for *Chac1* in sorted Beta-cell enriched fraction. (d) Each dot represents a pooled sample of 5 islets from one mouse, 3 mice were used per experiment, (e) one dot represents a 10 islet sample of a unique mouse. Statistics: two-sided Mann–Whitney U test; Error bars represent SD; ★★p < 0.01.

## Discussion

In the present study we explored the function of the cystine/glutamate transporter sXc *in vitro* as well as *in vivo* in the context of beta-cell and macrophage function.

At 12 weeks of age, sXc KO mice have a mildly increased glucose disposal. At 26 weeks of age, sXc-deficient mice additionally develop mildly lower insulin secretion. This suggests that the observed insulin secretory phenotype is relatively subtle and takes longer to manifest. Additionally, there seems to be a compensatory increase in glucose disposal in response to lower insulin secretion resulting in lower or unchanged glycemia. This could be a compensatory response to constitutive lower insulin secretion, increasing insulin sensitivity. This could also suggest immune cells deficient in *Slc7a11* might not be able to induce insulin resistance as effectively. The observed lower insulin secretion is not caused by lower beta-cell mass. This suggests that there might be a reduction in functional secretory capacity of the beta-cells or an innate increase in insulin sensitivity.

When assessing insulin secretion by *ex vivo* glucose-stimulated insulin secretion (GSIS), we found lower insulin output in Slc7a11 KO islets, corroborating our *in vivo* findings and supporting an islet intrinsic defect over a systemic adaptation. Addition of the glutathione precursor γ-Glutamyl-Cysteine (GGC) abolished the difference in insulin secretion between WT and KO islets. It has to be noted however, that overall insulin output was significantly decreased in both groups by the treatment. This could be explained by endoplasmic reticulum stress through disruption of the organelle’s redox potential or GGC’s anti-oxidative properties and its ability to act as a co-factor for glutathione peroxidases. GGC could scavenge the ROS normally produced upon high-glucose stimulation, that couple glucose metabolism to insulin secretion.

Lower glutathione could also contribute to a lack of redox buffering capacity within the cell, namely through glutathione’s role as co-factor in ROS-detoxifying enzymes. We were not able to confirm this hypothesis using flow cytometry-based redox measurements in dispersed islets. However, these dyes are not very sensitive, so it is possible that sXc deficient beta-cells are experiencing low-grade redox stress that can be compensated for and that is too subtle to measure with the currently employed methods. Further, lower glutathione levels might also affect mitochondrial physiology. Indeed, we found lower levels of ATP in the islets, which could be either attributed to lowered cell viability or diminished mitochondrial metabolism.

The differential glucose metabolism observed in chow-fed animals disappeared when mice were fed a HFD. Slc7a11KO animals in this setting were not able to increase their insulin secretion to similar levels as WT animals. Despite the islet expansion induced by HFD and lower insulin secretion in response to a glucose bolus, there was no difference in beta-cell mass between genotypes, again indicating a functional defect not related to beta-cell maintenance or expansion.

*Chac1* was upregulated in sorted beta-cells and the CD45+ immune cell fraction. CHAC1 is a glutathione-degrading enzyme upregulated upon cystine starvation. This could therefore be a factor further driving down the islet glutathione levels, suggesting glutathione pools can be readily depleted to prioritize other uses for cysteine, as a mechanism to mitigate cysteine starvation. When we sought to monitor the effect of sXc deficiency on thiol levels, we found no difference in total thiol levels in islet cell fractions. Despite the lower glutathione levels, the thiol levels in islets were comparable between genotypes. This suggests that, despite lower glutathione, the total thiol levels of sXc deficient islet cells are similar between genotypes. Lacking the transporter is therefore compensated for by higher thiol retention in islet cells or compensatory upregulation of other pathways through which the cell can acquire sulphur containing metabolites.

The high Slc7a11 expression levels of islet immune cells compared to other islet fractions prompted us to study the myeloid-specific role of Slc7a11 in islet function. At 26 weeks of age, we did not find a similar phenotype as in constitutive Slc7a11 knockout mice. Additionally, islets isolated from these myeloid cell-specific Slc7a11 knockout mice did not have lower glutathione levels, nor did they have decreased GSIS. This suggests that the insulin secretory phenotype is mostly mediated by sXc deficiency in non-myeloid islet cells, and that the increased glucose disposal might be secondary to lower insulin secretion.

When fed a HFD for 4 weeks, insulin secretion trended to be lower in the KO animals but this failed to reach statistical significance. Longer HFD feeding dissipated the metabolic phenotype. This fits a common pattern in mouse metabolic phenotypes that are either severe enough to completely decompensate over time or mild enough to allow the metabolic system to adapt to the initial disruption and ultimately compensate the homeostatic disturbance. This could mean that sXc’s role is most prominent in the initial phase of adaptation to metabolic perturbation by dietary intervention.

Beta-cell mass was similar in control and tissue-specific KO in both males and females, excluding a role of myeloid cell sXc in islet expansion and maintenance. This is of importance, because excessive glutamate as observed in beta-cell lines exposed to high glucose concentrations, promotes apoptosis, which is mediated by N-methyl-D-aspartate (NMDA) receptor signalling. NMDA receptor inhibition has also been shown to increase the depolarization plateaus observed during insulin secretion, leading to higher insulin output and more efficient glucose disposal. Furthermore, sXc on astrocytes and microglia has been implicated as the main culprit in glutamate-induced neurotoxicity[18](26)[29]. We were not able to find an improvement in islet function upon myeloid sXc deletion. These data therefore suggest that myeloid sXc does not contribute to glutamate induced excitoxicity of beta-cells.

Concluding, sXc whole body deficiency leads to impaired insulin secretion and a mild cysteine shortage as evidenced by *Chac1* upregulation. Myeloid-specific sXc disruption does not affect insulin secretion or cysteine shortage, and does not protect against HFD-induced islet dysfunction.

## Acknowledgments

The authors thank Hélène Mereau for excellent technical assistance.

## Author Contributions

AB designed the study. AB and LR performed and analyzed experiments. AB, DTM, MBS and MYD wrote the manuscript. AF conceived the original idea. MYD is the guarantor of this work and, as such, had full access to all the data in the study and takes responsibility for the integrity of the data and the accuracy of the data analysis.

