## Supplemental Figures for "Cystine/glutamate antiporter system Xc^-^ deficiency impairs insulin secretion"

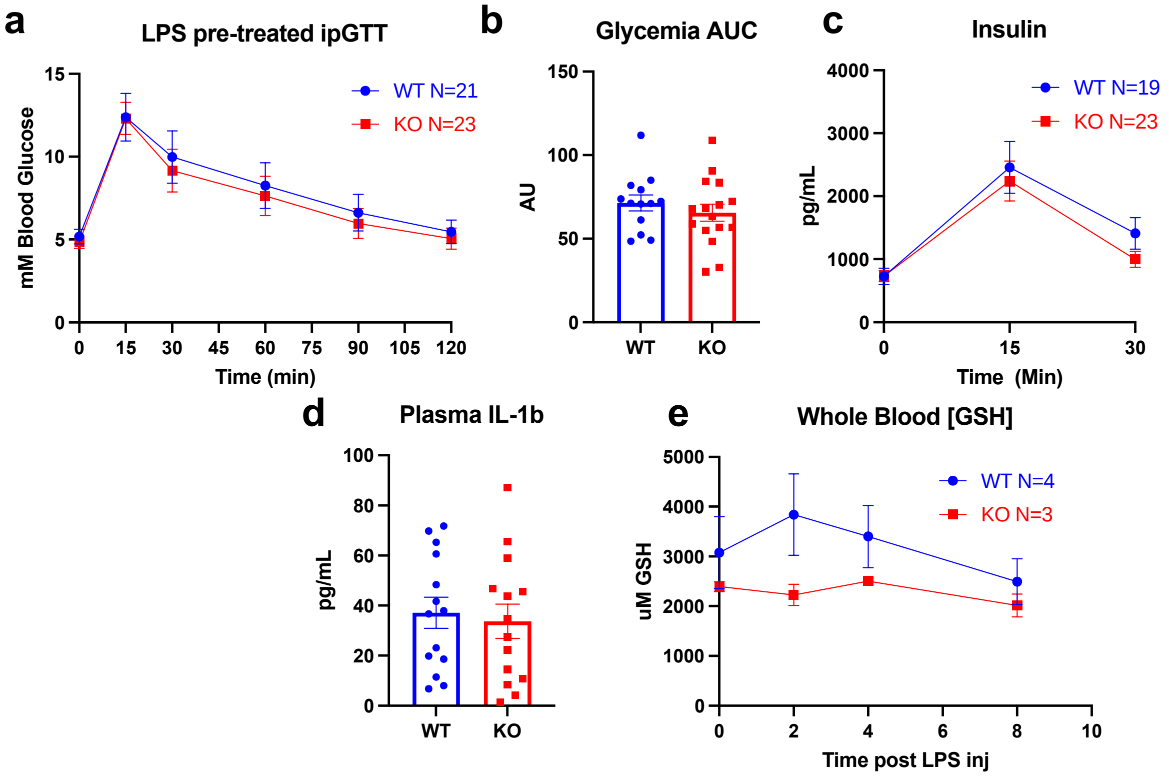


Figure S1 ipGTT with LPS pre-treatment 6 hours before experiment in whole body Slc7a11 deficient animals, (a) ipGTT glycemia, (b) corresponding AUC, (c) circulating insulin and after intraperitoneal injection of 2g/kg glucose following a 6h fast, (d) plasma IL-1b 6 hours post LPS injection, (e) whole blood glutathione timecourse after 2mg/kgbw LPS injection. Error bars represent SD.

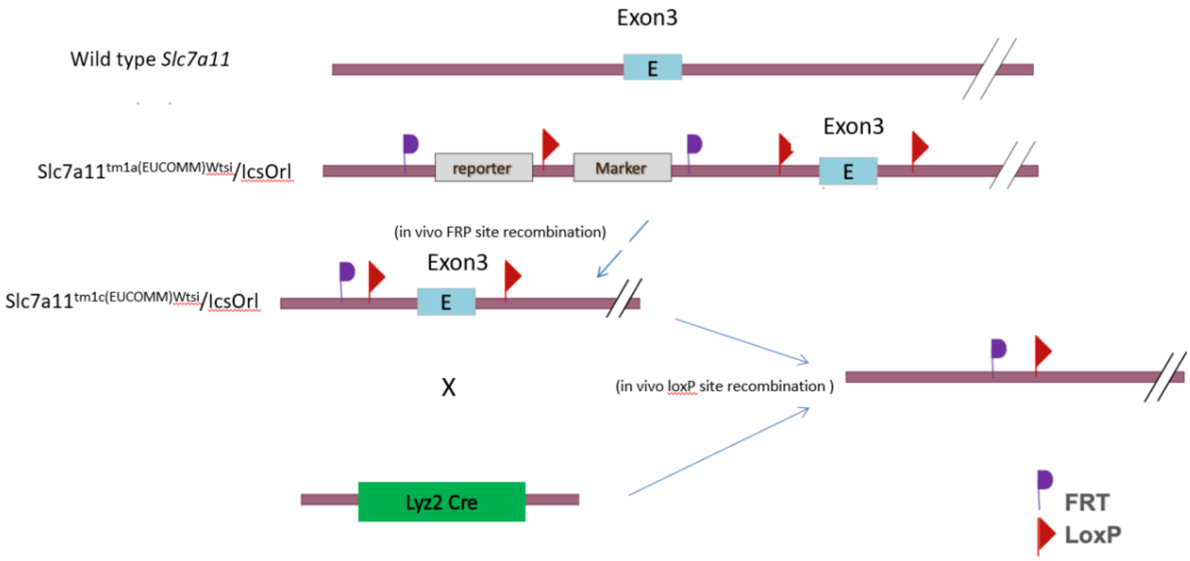


Figure S2 Recombinations strategy for generation the myeloid specific *Slc7a11* deficient animals.The mice carrying the tm1a allele were crossed with a flipase transgenic mouse to remove the FRT-flanked report and marker cassette. Subsequently, by crossing homozygous tm1c carrying animals with a Cre-recombinase driven by a Lyz2 promoter, myeloid specific *Slc7a11* deficiency was established.

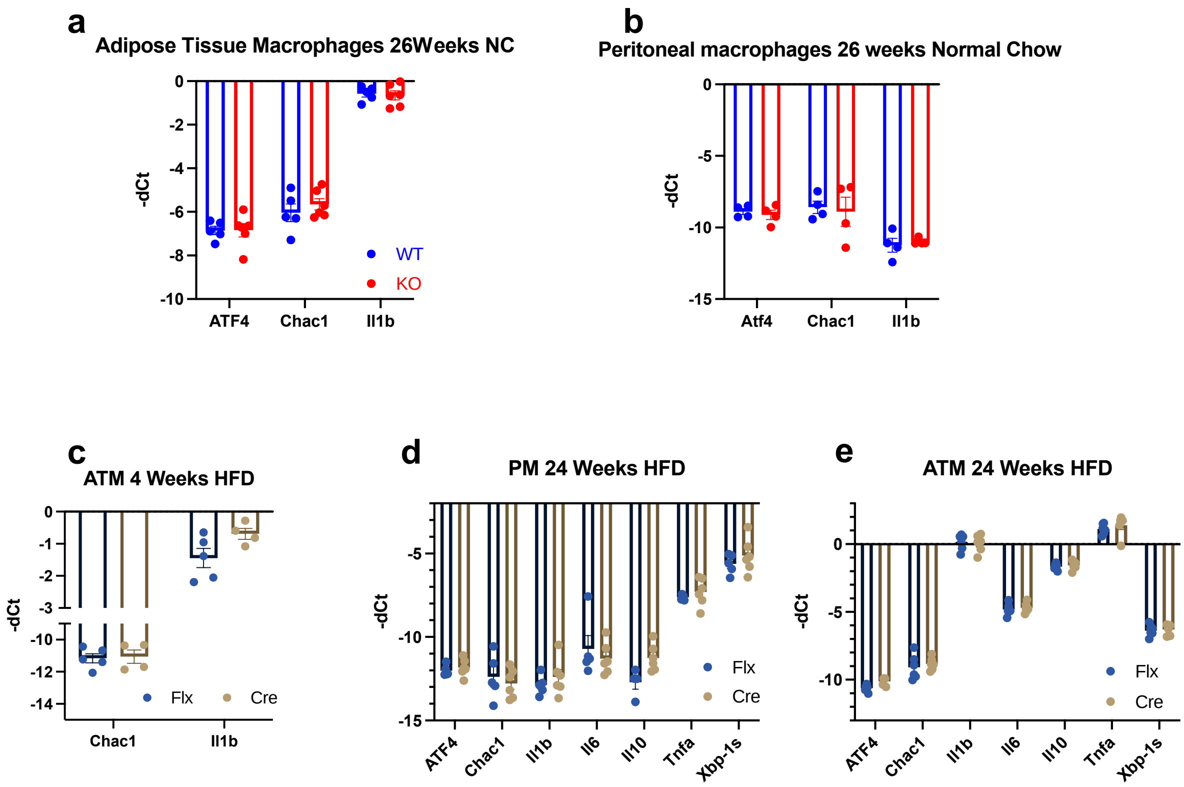


Figure S3 qPCR screening of sorted CD11b+ F480+ macrophages from adipose tissue and peritoneal lavage: Whole body KO: (a,b) qPCR of peritoneal and adipose tissue macrophages from mice 26 weeks old mice.

Myeloid specific KO: (c) rtPCR of ATMs from mice fed 4 weeks HFD (d,e) peritoneal and adipose tissue macrophages from 24W HFD fed male mice.

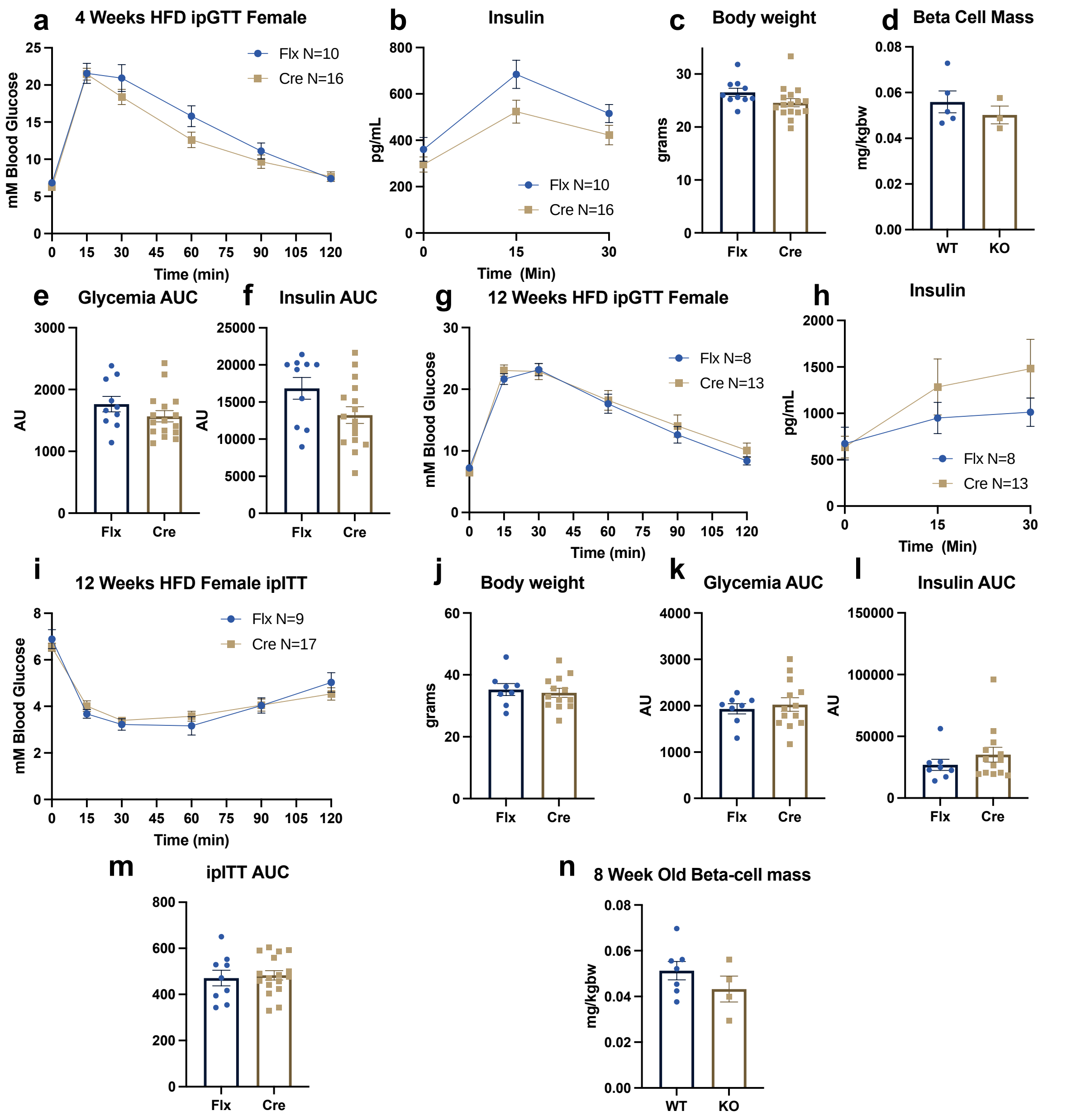


Figure S4 LysM Cre Slc7a11 Flx Females metabolic experiments
(a,b) ipGTT, (c) body weight, (d) Beta-cell mass, (e,f) area under the curve of a&b respectively, of mice fed HFD for 4 weeks, (g,h,j,k,l) ipGTT and body weight of mice fed HFD for 12 weeks, (i,m) ITT of mice fed HFD for 12 weeks, (n) Beta-cell mass of 8 week old mice (before start HFD). Statistics: two-sided Mann–Whitney U test; error bars represent SD.

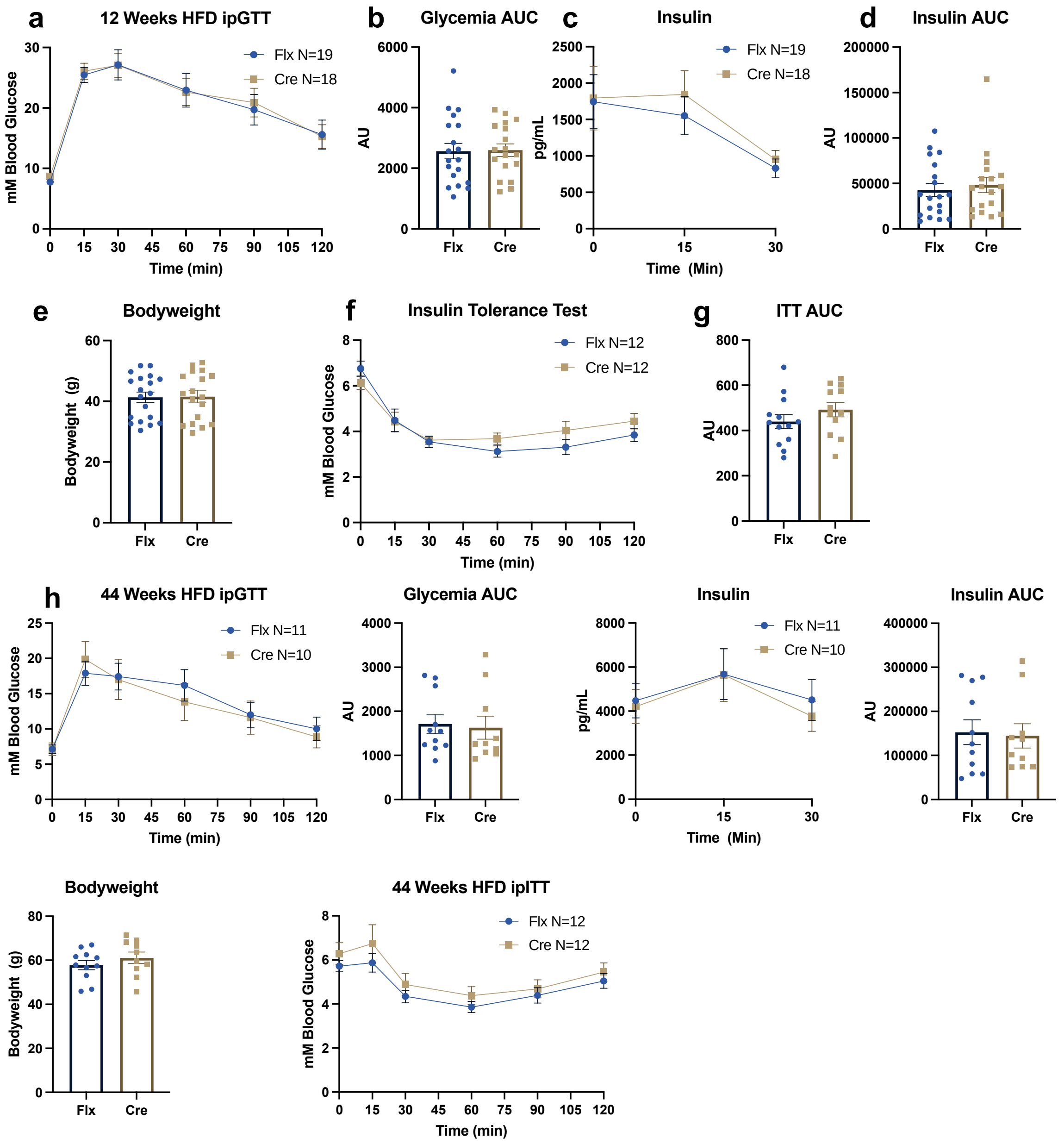


Figure S5 LysM Cre Slc7a11 Flx fed HFD for 12 weeks (a) ipGTT glycemia, (b) corresponding AUC, (c) circulating Insulin and (d) corresponding AUC after intraperitoneal injection of 2g/kg glucose following a 6h fast, (e) body weight, (f,g) insulin tolerance test. LysM Cre Slc7a11 Flx fed HFD for 44 weeks (h) ipGTT glycemia, (i) Insulin, (j) AUC of glycemia, (k) AUC of insulin, (l) Bodyweight ,(m,n) insulin tolerance test. Error bars represent SD.
